# Modeling small RNA competition in *C. elegans*

**DOI:** 10.1101/021576

**Authors:** Joshua M. Elkington

## Abstract

**Website Summary:** Small RNAs are important regulators of gene expression; however, the relationship between small RNAs is poorly understood. Studying the crosstalk between small RNA pathways can help understand how gene expression is regulated. One hypothesis suggests that small RNA competition arises from limited enzymatic resources. Therefore, a model was created in order to gain insights into this competition.

**Abstract:** Small RNAs have been determined to have an essential role in gene regulation. However, competition between small RNAs is a poorly understood aspect of small RNA dynamics. Recent evidence has suggested that competition between small RNA pathways arises from a scarcity of common resources essential for small RNA activity. In order to understand how competition affects small RNAs in *C. elegans*, a system of differential equations was used. The model recreates normal behavior of small RNAs and uses random sampling in order to determine the coefficients of competition for each small RNA class. The model includes endogenous small-interfering RNAs (endo-siRNA), exogenous small-interfering RNAs (exo-siRNA), and microRNAs (miRNA). The model predicts that exo-siRNAs is dominated by competition between endo-siRNAs and miRNAs. Furthermore, the model predicts that competition is required for normal levels of endogenous small RNAs to be maintained. Although the model makes several assumptions about cell dynamics, the model is still useful in order to understand competition between small RNA pathways.

## Introduction

In *C. elegans* there are three main classes of small RNAs: endogenous small-interfering RNAs (endo-siRNA), microRNAs (miRNA) and piwi-interacting RNAs (piRNA). Furthermore, exogenous dsRNA can be delivered to worms either by feeding or injection in order to induced RNA interference (RNAi). The dsRNA delivered to worms are processed by proteins such as Dicer in order to produced siRNAs that associate with proteins such as Argonautes in order to produce a RNA-induced silencing complex (RISC) that silences target mRNAs by degradation if there is perfect base pairing between the siRNA and the mRNA (Duchaine, 2006) (Figure 1a). C. elegans possess a single Dicer protein (DCR-1) (Knight, 2001) that also processes precursor miRNAs to create mature miRNAs (Ketting, 2001) and that helps create endo-siRNAs. miRNAs regulate gene expression in similar way as exo-siRNAs and endo-siRNAs, but miRNAs base-pairing to the 3’ untranslated region (UTR) of the target mRNAs to degrade the target of inhibit translation depending on the degree of complementary with the target (Lanford, 2010). miRNAs were discovered in *C. elegans*; however, small RNAs have been found to be broadly conserved in animals and plants (Aalto, 2012). Small RNAs are known to regulate germline maintenance (Han, 2009), viral protection (Knight, 2001), development, RNAi inheritance, and chromosomal segregation (Ketting, 2001) (Figure 1b).

**Figure 1.**
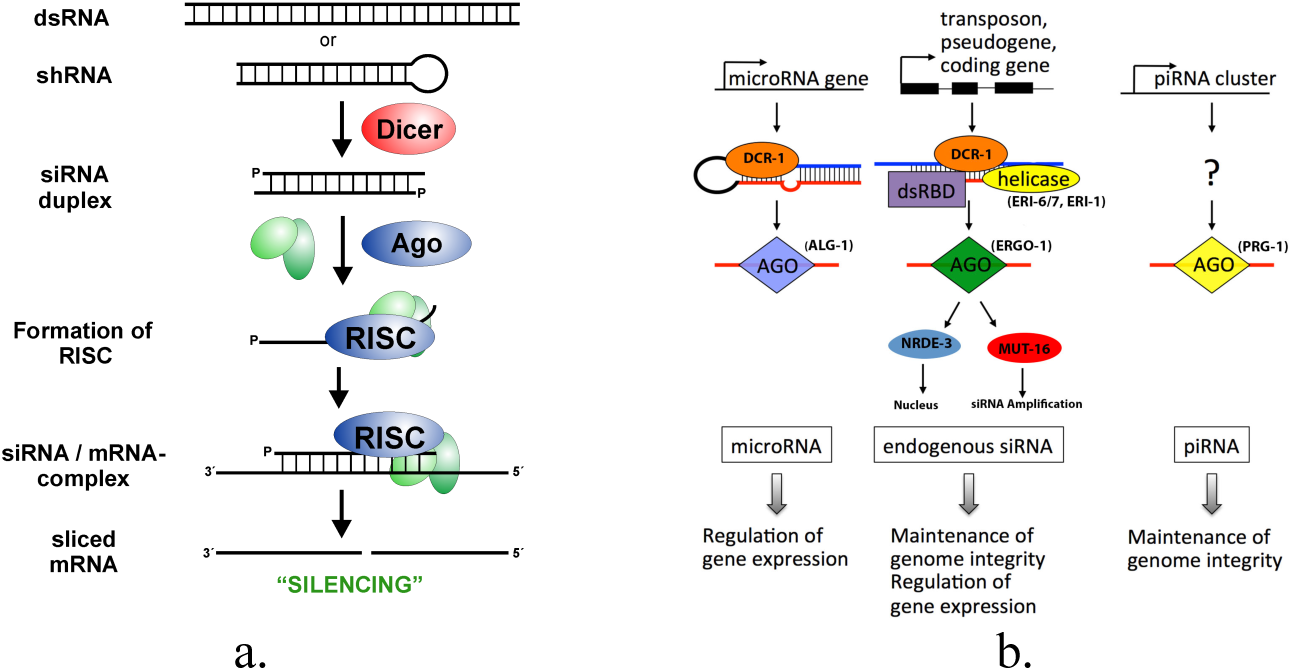
Outline of RNAi and Small RNA pathways a. Processing of dsRNA or shRNA into mature siRNAs within a RISC by Dicer and Ago proteins (http://www.gene-quantification.de/rnai.html) b. Outline of three endogenous small RNA pathways with function in *C. elegans*.

Recently there have been findings suggesting that small RNA pathways compete for common resources within *C. elegans*. Disrupting endogenous small-interfering RNAs (endo-siRNA) enhances *C. elegans* sensitivity to exogenous RNA interference (exo-RNAi) and microRNA (miRNA) efficacy (Zhuang, 2012). Mutant *C. elegans* sensitive to exo-RNAi were found to have lower levels of endo-siRNAs (Vasale, 2010 & Simmer, 2002). These small RNAs use Dicer and Argonaute proteins to function properly (Guang, 2010), and competition may arise from the limited amount for these proteins. Moreover, the competition between small RNA pathways may also included RNA-dependent RNA polymerases (RdRP) and secondary siRNA binding Argonautes (SAGO). Overexpression of SAGO-1 and SAGO-2 causes enhanced exo-RNAi in worms (Yigit, 2006). The loss of miRNAs 35-41 in *C. elegans* led to hypersensitivity to exo-RNAi and lower levels of endo-siRNAs (Massirer, 2012).

Despite recent evidence implicating competition between exo-RNAi and endo-siRNAs, there has been little evidence showing competition with other small RNA pathways. In *C. elegans*, miRNAs are predicted to regulate more than half of the transcriptome; therefore, competition between small RNA pathways such as miRNAs and endo-siRNAs is important to understand the regulation of gene expression. So understanding cross-regulation between exo-RNAi, endo-siRNA, and miRNA may help elucidate several important pathways such as development and viral protection.

In order to gain a better understanding of the competition between small RNA pathways, a system of differential equations was created to model this competition. The system modeled the behavior of exo-siRNAs, endo-siRNAs, and miRNAs in order to determine the competition coefficients between the small RNA classes. Furthermore, piRNAs were later included to further refine the model. This math model is useful to understand the competition for resources between different types of small RNAs. Finding a quantitative value for the effects of competition on small RNAs can offer insights into the dynamics of endogenous small RNAs and the effects of RNAi on other small RNA pathways. The model can be scaled to include other small RNA pathways in order to increase accuracy and gain more information on small RNA competition.

## Methods

In order to create a realistic model of small RNA competition, production rates and protein carrying capacity were determined by measured values. The competition coefficients were determined by random sampling to obtain the observed behavior of small RNAs. The model assumes that exogenous dsRNA delivered to worms induces an RNAi effect that eventually decreases over time. The production rate of the three small RNA classes is proportional to the levels of Dicer and Argonaute proteins. This proportionality was modeled using the Monad model, which uses some parts of the Michaelis-Menten equation. Furthermore, the model assumes that Dicer and Argonaute proteins grow logistically with a carrying capacity.

For initial conditions, the model assumes all proteins and small RNAs have an initial concentration of 500 nM except for exogenous small RNAs. The model assumes that due to injection or feeding that exogenous small RNAs over saturate the cell. The production rates were determined from molecular half-life using the following equations: 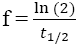 and N(t) = N*e*^*−ft*^. I assumed that protein half-life is 46 hours and protein number per cell is 50,000 (Schwanhäusser, 2011). I assumed miRNA half-life is 119 hours (Gantier, 2011), exo-RNAi half-life is 48 hours (Chiu, 2003), and endo-siRNA half-life is 72 hours (Bartlett, 2006). The average volume of a C. elegans cell is approximately 50 *mm*^3^, and the protein carrying capacity was derived from this volume and the average protein number.

## Results

After replicating the desired behavior of the small RNA classes (Figure 2), the model predicts the competition from miRNA and endo-siRNAs dominate exo-siRNAs. However, miRNA levels take a relatively longer time to recover (Figure 2c), which is likely due to competition effect from endo-siRNAs. For the model behavior to be created, miRNA or endo-siRNA must “win” over the other. Moreover, when exo-RNAi was removed from the model, endo-siRNAs grow linearly and miRNA oscillates to zero. When endo-siRNAs or miRNAs were removed, the other small RNA was still able to outcompete exo-siRNAs.

**Figure 2.**
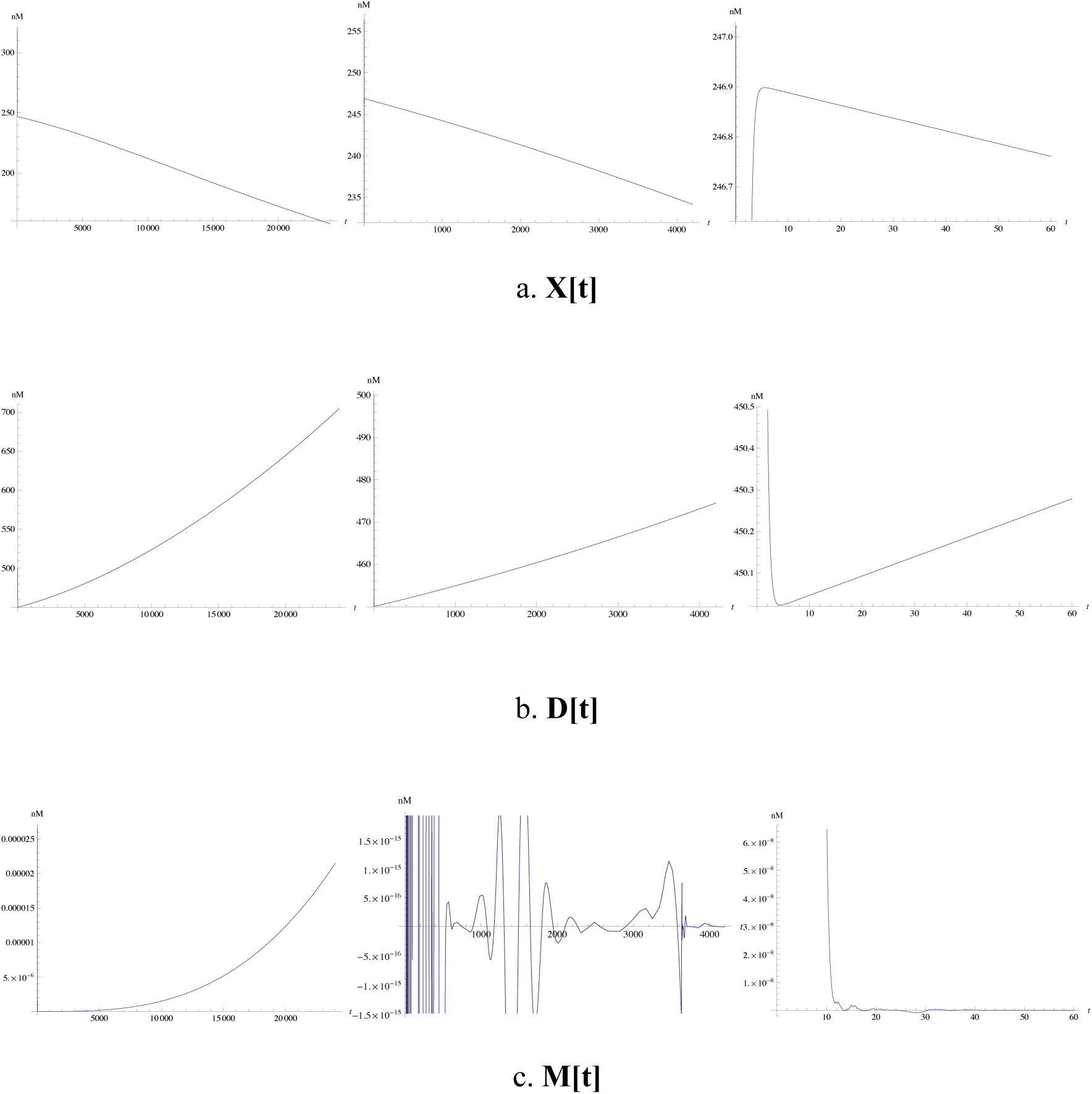
Small RNA behavior. The y-axis of the graphs is concentration in nM and the x-axis is time in minutes. The leftmost graphs are over 24,000 minutes, the middle graphs are over 4,200 minutes, and the rightmost graphs are over 60 minutes. a. Concentration of exo-siRNAs with respect to time. b. Concentration of endo-siRNAs with respect to time. Concentration of miRNAs with respect to time.

In order to create a more accurate model to understand competition between various small RNA pathways, piRNAs were included in the model. The model assumes that piRNAs only compete for Argonaute proteins because they do not use Dicer. With piRNAs included without exo-RNAi, piRNA levels initially decrease but recover over time (Figure 3). Furthermore, miRNA levels were able to increase with respect to time even without exo-RNAi. The modified model predicts that the effect of competition between piRNAs and miRNAs is the same for both. The competition effect of piRNAs from endo-siRNAs is more significant than competition on endo-siRNAs from piRNAs. When exo-RNAi was included in this modified model, the same behavior from the previous model (Figure 2) is observed. In this model, exo-siRNAs are dominated by competition by piRNAs.

**Figure 3.**
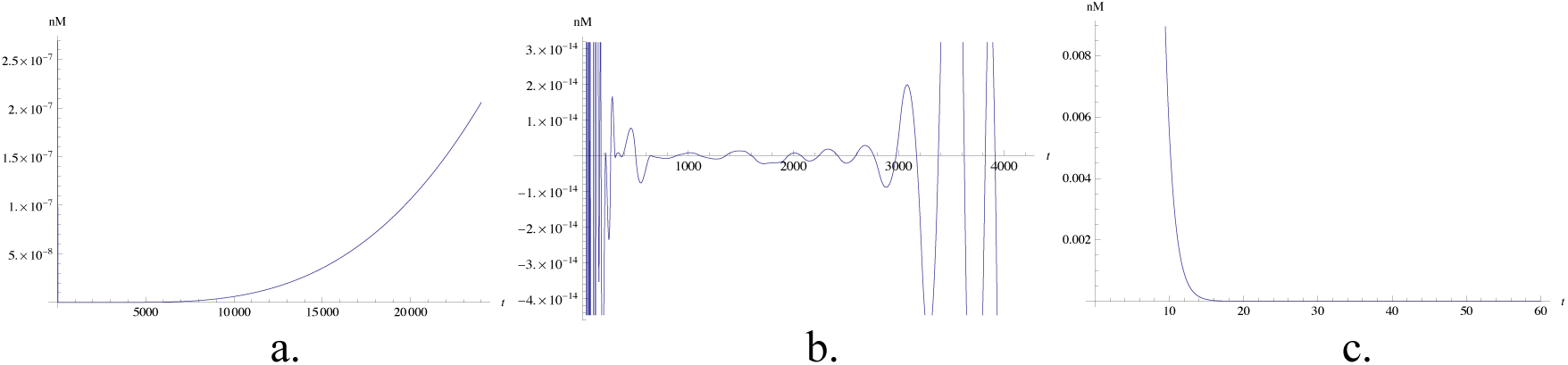
piRNA concentration without RNAi. a. Concentration (nM) of piRNAs over 24,000 minutes. b. Concentration (nM) of piRNAs over 4,200 minutes. c. Concentration (nM) of piRNAs over 60 minutes.

**Figure 4.**
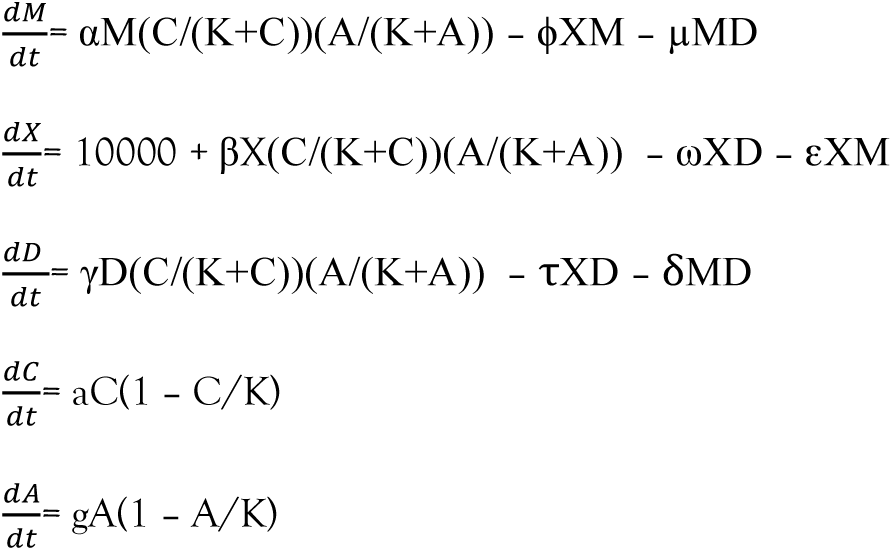
Small RNA Competition Model. System of differential equations used to model competition between miRNAs, endo-siRNAs, and exo-siRNAs within *C. elegans*.

## Conclusion

The model makes predictions for the competition between small RNAs. Within the model, exo-siRNAs “lose” the competition between endogenous small RNAs. As a result, this competition may cause exo-RNAi to lose efficacy over time. Without exo-RNAi, miRNA levels oscillate to zero, which predicts the miRNA activity is high in the beginning of the cell life then overtime miRNA activity is silenced. So there may be other molecules that compete with miRNAs and endo-siRNAs for resources because this is not the observed behavior of miRNAs. Therefore, piRNAs were included in the model to understand small RNA competition without exo-RNAi. Without exo-RNAi, the miRNAs were able to increase over time when piRNAs were included. Competition between small RNA pathways may be needed for growth of small RNAs because the checks and balances don’t allow one small RNA class to “win.”

This model implies that competition between various small RNA pathways varies across classes. So the effect of small RNA competition may arise from cellular localization, concentration, and levels of resources. Ultimately, the model can be made more accurate by modeling resources individually. So each Argonaute, RdRP, and other small RNA related proteins can be included within the model (Figure 1b). Furthermore, the half-life of each of these proteins can be experimentally determined in order to achieve better results. The model can be scaled up or down to study broad classes of small RNAs or study individual small RNAs if some of the values of found experimentally. Understanding small RNA competition is important to gain insights into the effect of exo-RNAi and also understand endogenous small RNA dynamics.

---

**Table 1.** Terms. Overview of terms used in system of differential equations.

| Terms |  |
| --- | --- |
| $M(t)$ | miRNA (nM) |
| $X(t)$ | exo-siRNA (nM) |
| $D(t)$ | endo-siRNA (nM) |
| $C(t)$ | Dicer (nM) |
| $A(t)$ | Argonaute (nM) |
| $\alpha$ | production rate of miRNA |
| $\beta$ | production rate of exo-siRNA |
| $\gamma$ | production rate of endo-siRNA |
| $\phi, \varepsilon$ | competition rate between $M(t)$ and $X(t)$ |
| $\omega, \tau$ | competition rate between $D(t)$ and $X(t)$ |
| $\mu, \delta$ | competition rate between $D(t)$ and $M(t)$ |
| $a$ | production rate of $C(t)$ |
| $g$ | production rate of $A(t)$ |
| $K$ | carrying capacity of $C(t)$ and $A(t)$ |
| $t$ | time (minutes) |

**Table 2.** Term Values. Term values determined from previous experimental data or obtained by random sampling.

| Term Values |  |
| --- | --- |
| $\alpha$ | 0.0000339 |
| $\beta$ | 0.0000926 |
| $\gamma$ | 0.0000926 |
| $\phi$ | 0.00000000005 |
| $\varepsilon$ | 0.09 |
| $\omega$ | 0.09 |
| $\tau$ | 0.00000000005 |
| $\mu$ | 0.005 |
| $\delta$ | 0.0005 |
| $a$ | 0.000227 |
| $g$ | 0.0001135 |
| $K$ | 1000 nM |
| $M(0)$ | 500 nM |
| $X(0)$ | 10,000 nM |
| $D(0)$ | 500 nM |
| $C(0)$ | 500 nM |
| $A(0)$ | 500 nM |

